# Cancer cells are sensitive to wild-type IDH1 inhibition under nutrient limitation

**DOI:** 10.1101/2020.11.19.390633

**Authors:** Ali Vaziri-Gohar, Jonathan J. Hue, Hallie J. Graor, Erin Prendergast, Vanessa Chen, Joel Cassel, Farheen S. Mohammed, Ata Abbas, Katerina Dukleska, Imran Khokhar, Omid Hajhassani, Mahsa Zarei, Rui Wang, Luke D. Rothermel, Ilya Bederman, Jessica Browers, Robert Getts, Henri Brunengraber, Joseph M. Salvino, Jonathan R. Brody, Jordan M. Winter

## Abstract

Pancreatic cancer cells alter their metabolism to survive cancer-associated stress (1-4). For example, cancer cells must adapt to steep nutrient gradients that characterize the natural tumor microenvironment (TME) (5-7). In the absence of adaptive strategies, harsh metabolic conditions promote the generation of free radicals (8) and impair energy production in tumor cells. Towards this end, wild-type isocitrate dehydrogenase 1 (IDH1) activity is a metabolic requirement for cancer cells living in a harsh metabolic milieu. The cytosolic enzyme interconverts isocitrate and alpha-ketoglutarate, and uses NADP(H) as a cofactor. We show that under low nutrient conditions, the enzymatic reaction favors oxidative decarboxylation to yield NADPH and alpha-ketoglutarate. Metabolic studies showed that the IDH1 products directly support antioxidant defense and mitochondrial function in stressed cancer cells. Genetic IDH1 suppression reduced growth of pancreatic cancer cells in vitro under low nutrient conditions and in mouse models of pancreatic cancer. Surprisingly, allosteric inhibitors of mutant IDH1 proved to be potent wild-type IDH1 inhibitors under conditions specific to the TME, highlighting a natural therapeutic window. The presence of low magnesium enhanced allosteric inhibition by the drug, and ambient low glucose levels enhanced cancer cells’ dependence on wild-type IDH1. Thus, intrinsic TME conditions sensitized wild-type IDH1 to FDA-approved AG-120 (ivosidenib), and revealed the drug to be a potent single-agent therapeutic in cell culture and diverse in vivo cancer models. This work identified a potentially new repertoire of safe cancer therapies, including a clinically available compound, for the treatment of multiple wild-type IDH1 cancers (e.g., pancreatic).

---

Pancreatic cancer (PC) is the third leading cause of cancer death in the United States (*9*). Progress in clinical outcomes is largely attributable to modest survival gains observed with multi-agent chemotherapy combinations. Our recent work revealed that harsh conditions in the PC microenvironment actually induce chemo-resistance (*4*). Thus, the best available treatments (i.e., chemotherapy) fail to target biologic vulnerabilities. A more compelling and strategic therapeutic approach should identify metabolic dependencies in the context of austere conditions that characterize primary and metastatic PC (*6, 7, 10*), as well as other cancers (*11–14*). Examples include nutrient scavenging mechanisms like autophagy (*15*) and macropinocytosis (*7, 16*).

Glucose and glutamine fuel ATP and NADPH synthesis pathways. These biochemical products comprise the basic energy and reductive currencies in cancer cells. Thus, under nutrient-deprived conditions, cancer cells benefit from adaptive strategies that maximize ATP generation (*17*) and neutralize free radicals (*18*). To validate this premise, PC cells were cultured under glucose concentrations (2.5 mM) half that of serum and normal tissues (Extended Data Fig. 1a) to simulate levels in the PC microenvironment (*7*). A surge in reactive oxygen species (ROS) levels (Fig. 1a) occurred over 48 hours, reflecting the oxidative stress caused by glucose withdrawal. Cells compensated and levels returned to baseline the following day. Despite low ambient glucose levels, a compensatory rise in NADPH occurred (Fig. 1b, Extended Data Fig. 1b) to power the observed ROS neutralization. An siRNA screen of 13 NADPH-generating enzymes was performed to investigate a possible mechanism behind the compensatory response. IDH1 silencing reproducibly lowered reductive power under glucose withdrawal (2.5 mM for 72 hours) in two different PC cell lines (MTT assay, Fig. 1c, Extended Data Fig. 1c). ELAVL1 (HuR) silencing was used as a positive control (*4, 19*). *IDH1* mRNA expression reproducibly increased under glucose withdrawal across PC cell lines (Fig. 1d), supporting a prior finding that the enzyme is part of an acute survival response to nutrient withdrawal (*4, 19*).

**Figure 1.**
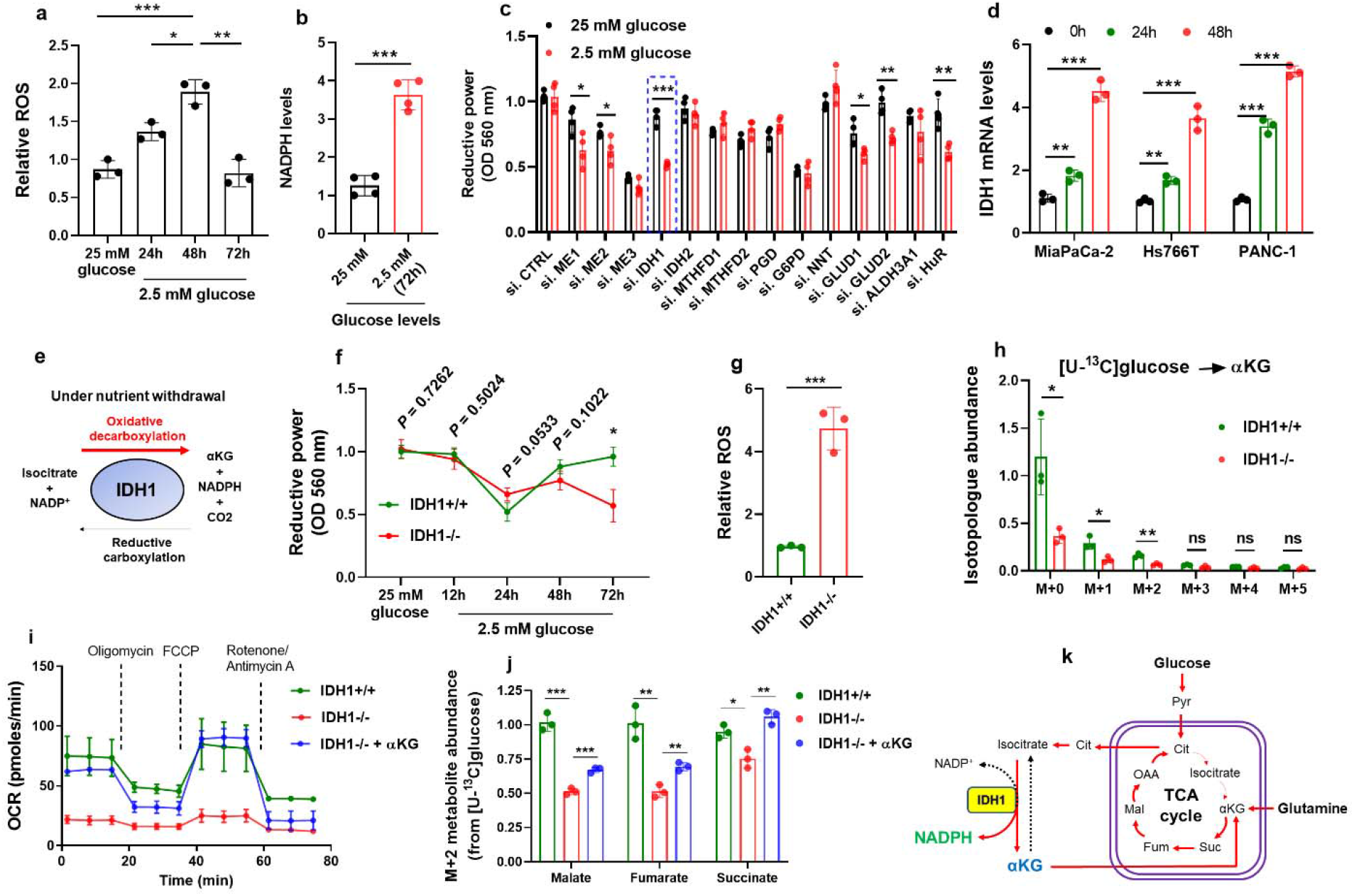
wtIDH1 supports antioxidant defense and mitochondrial function under nutrient withdrawal in pancreatic cancer cells. **a**, ROS levels were detected by DCFDA assay in MiaPaCa-2 pancreatic cancer (PC) cells under glucose withdrawal (2.5 mM) over 72 hours. **b**, NADPH levels in MiaPaCa-2 PC cells under the indicated conditions for 72 hours; n=4 independent experiments. **c**, Reductive power, as detected by an MTT assay, normalized to cell number, in MiaPaCa-2 PC cells. Cells were transiently transfected with siRNAs against different NADPH-generating enzymes and incubated under the indicated conditions for 72 hours. HuR silencing was performed as a positive control; n=4 independent experiments. **d**, qPCR analysis of *IDH1* mRNA normalized to 18S in multiple PC cells and under 2.5 mM glucose for the indicated time interval. **e**, Enzymatic reaction of IDH1. **f**, Reductive power in MiaPaCa-2 PC cells under nutrient withdrawal over 72 hours. **g**, ROS levels were detected 72 hours after incubation of MiaPaCa-2 PC cells under glucose withdrawal. **h**, Representative GC-MS analysis of glucose-derived αKG, normalized to M+0 from IDH1+/+ cells, in MiaPaCa-2 PC cells under glucose withdrawal; n=2 independent experiments. **i**, Representative oxygen consumption rate (OCR) in MiaPaCa-2 PC cells treated initially with αKG (4 mM) for 6 hours followed by culturing cells in 2.5 mM glucose for 30 hours; n=2 independent experiments. **j**, Representative GC-MS analysis of relative abundance of glucose-derived M+2 metabolite enrichment in MiaPaCa-2 PC cells after treatment with αKG (4 mM); n=2 independent experiments. **k**, Schematic illustration of IDH1-derived αKG anaplerosis into the TCA cycle. Pyr, pyruvate; Cit, citrate; Suc, succinate; Fum, fumarate; Mal, malate; OAA, oxaloacetate. Data are provided as mean ± s.d. from three independent experiments, unless indicated. *P* values were calculated using two-tailed, unpaired Student’s *t*-tests. *, P < 0.05; **, P < 0.01; ***, P < 0.001.

Prior studies in melanoma and lung cancer cells first identified a possible role for wild-type (wt) IDH1 in cancer cell biology (*20–22*). These studies demonstrated that NADPH-driven reductive carboxylation catalyzed by wtIDH1 promoted lipid synthesis in proliferating cancer cells. A much greater body of literature has focused on the metabolic implications of neomorphic mutant (mt) IDH1. As with reductive carboxylation of the wtIDH1 isoenzyme, mtIDH1 consumes NADPH and _α_-ketoglutarate (αKG), but generates 2-hydroxyglutarate instead of isocitrate (*23*). We hypothesized that the alternative wtIDH1 reaction, oxidative decarboxylation, is favored under low nutrient conditions. NADPH and αKG produced by this reaction could support antioxidant defense (primarily through NADPH synthesis) and mitochondrial function (primarily through anaplerosis of αKG) to promote cancer cell survival when nutrients are scarce (Fig. 1e).

IDH1-knockout PC cell lines (IDH1−/−) were generated via CRISPR-Cas9 gene editing and validated by Western blot and sequencing (Extended Data Fig. 1d). *IDH2* and *IDH3* expression were similar between IDH1−/− and isogenic controls (IDH1+/+) (Extended Data Fig. 1e). Initial studies tested the effects of IDH1 deletion on redox homeostasis. As observed with parental PC cells (Fig. 1a), IDH1+/+ cells adapted to oxidative stress from glucose withdrawal. In contrast, IDH1−/− cells failed to compensate (Fig. 1f, Extended Data Fig. 1f). Impaired redox balance in IDH1−/− cells was corroborated in a DCFDA assay, which showed increased ROS levels with glucose withdrawal (Fig. 1g). IDH1−/− cells cultured under low glucose conditions were also sensitive to additional oxidative insults, like hydrogen peroxide (Extended Data Fig. 1g) or chemotherapy (Extended Data Fig. 1h). In the latter experiment, stable re-expression of wtIDH1 rescued IDH1−/− PC cells (i.e., promoted chemotherapy resistance), while the catalytically-altered mtIDH1 did not. This finding validates a prior study by our group testing the growth of an IDH1-mutant cell line in a xenograft model. After forced suppression of both isoenzymes (mutant and wild-type) through CRISPR-deletion of a regulatory molecule, HuR, stable reconstitution of wtIDH1 had a far greater effect on tumor proliferation mtIDH1 expression (*24*). Although, IDH1 is localized to the cytoplasm, the effect of IDH1 deletion extended beyond this specific cellular compartment, mitochondrial-specific ROS dramatically increased (Extended Data Fig. 1i).

The impact of IDH1 on redox homeostasis tracks to changes in NADPH synthesis. The downstream consequences of altered _α_-ketoglutarate (αKG) levels (the other product of wtIDH1) are equally important through direct effects on mitochondrial metabolism. ^13^C metabolite tracing of uniformly-labeled ^13^C glucose revealed a marked decrease in αKG levels (M+2 fraction) in IDH1−/− PC cells cultured under low glucose conditions (Fig. 1h). This result confirms that oxidative decarboxylation (Fig. 1e) is favored under these conditions. We suspected that a reduction in IDH1-dependent αKG would negatively impact mitochondrial health, since αKG is an important anaplerotic substrate for the tricarboxylic acid (TCA) cycle (*22*). Consistent with others’ work (*17, 18*), parental PC cells (with normal levels of IDH1), adapt to glucose withdrawal by compensatorily increasing oxygen consumption and ATP production (Extended Data Fig. 2a, b). IDH1-knockout cells were not able to adapt (Fig. 1i and Extended Data Fig. 2c-e). αKG (4 mM) and Mito-Tempo (mitochondrial-specific antioxidant), rescued oxygen consumption capacity and ATP production in IDH1−/− cells cultured under low glucose conditions (Fig. 1i and Extended Data Fig. 2f), demonstrating the importance of this anaplerotic substrate for mitochondrial function. Interestingly, the difference in oxygen consumption between IDH1−/− and IDH1+/+ PC cells was negligible under nutrient replete conditions (25 mM glucose, Extended Data Fig. 2g). This observation highlights a recurring theme; wtIDH1 is unimportant for cell survival under nutrient-rich conditions.

The fate of αKG into the mitochondrial matrix and the TCA cycle was then mapped more precisely using additional ^13^C metabolites. While glucose-deprived IDH1−/− cells exhibited a reduction in carbon flux from [U-^13^C]glucose to αKG (due to the absence of wtIDH1, Fig. 1h), tracing with (1,2,3,4-^13^C_4_)αKG revealed that carbon flux from αKG, when present, is destined in part for the TCA cycle under these conditions (Extended Data Fig. 3a). This finding was further validated by tracing [U-^13^C]glucose into the TCA cycle. Under low glucose conditions, M+2 TCA metabolites were lower in IDH1−/− cells, as compared to IDH1+/+ cells (Fig. 1j). Again, the addition of αKG partially rescued IDH1−/− cells. In contrast to αKG, citrate (positioned upstream of the oxidative decarboxylation reaction) failed to stimulate carbon flux from [U-^13^C]glucose into the TCA cycle under low glucose conditions (Extended Data Fig. 3b, c). These experiments suggest that wtIDH1 is an important enzymatic node that influences the flow of carbon into and through the TCA cycle, as summarized in Fig. 1k. Carbon can move from the glycolytic pathway into the TCA cycle, exit via the citrate transporter, and re-enter the mitochondrial matrix as IDH1-derived αKG.

The impact of IDH1 on biochemical pathway flux also extends beyond the TCA cycle. Glucose uptake from the media into IDH1−/− PC cells increased, likely as an act of metabolic desperation in the face of scarce nutrients (Extended Data Fig. 3d). Despite increased glucose uptake, ^13^C-labeling of lactate from [U-^13^C]glucose was markedly diminished in IDH1−/− cells (Extended Data Fig. 3e). This finding was replicated through extracellular acidification rate (ECAR) measurements in multiple PC cell lines detected by extracellular flux analysis (Extended Data Fig. 3f, g). The reduction in lactate could not be explained merely by a reduction in M+2 lactate (from [U-^13^C]glucose) from the oxidative phase of the pentose phosphate pathway (PPP) (*25*) (Extended Data Fig. 3h). Rather, global dysfunction of metabolic pathways occurs in IDH1−/− cells. This observation was equally apparent in studies of glutamine utilization. Parental PC cells require glutamine under low glucose conditions, as evidenced by an increased nutrient uptake in parental PC cells (Extended Data Fig. 3i). However, ^13^C-flux into the TCA cycle from [U-^13^C]glutamine was markedly impaired in IDH1−/− PC cells (Extended Data Fig. 3j). Diminished ^13^C incorporation into lactate from [U-^13^C]glutamine was also observed in IDH1−/− cells (Extended Data Fig. 3k). Reduced carbon flux from glutamine into lactate suggests impaired anaplerosis of glutamine into the TCA cycle in IDH1−/− cells. These isotope tracer studies collectively suggest that IDH1-deletion point towards slower or less efficient TCA cycling (presumably due to the diminished anaplerotic flux of IDH1-derived αKG). Reduced glycine in IDH1−/− cells (Extended Data Fig. 3j) similarly suggests the impact of IDH1 in diverting carbon through metabolic pathways outside of the TCA cycle. Reduction in both glutamate and glycine levels may also conspire to hinder antioxidant defense, since these amino acids are principal components of the reducing agent, glutathione (Extended Data Fig. 3j). In light of the connection between wtIDH1 activity and glutamine metabolism, we tested the activity of a small molecule glutaminase inhibitor, CB-839, and observed increased sensitivity to the drug in IDH1−/− cells (Extended Data Fig. 3l). In addition to the effect on mitochondrial function, αKG also had an effect on ROS levels, as evidenced by a reduction in ROS levels with the addition of αKG in IDH1−/− cells (Extended Data Fig. 3m). The link between αKG and antioxidant defense may be attributable to the positive impact on mitochondrial efficiency (and consequently less ROS production) or support of other NADPH-generating enzymes, like malic enzyme (*26, 27*). These metabolic studies jointly suggest that wtIDH1 enhances antioxidant defense through the generation of NADPH and impacts metabolic pathways at the level of the TCA cycle through the generation of IDH1-derived αKG. Wild-type IDH1 also influences additional metabolic pathways outside of the TCA cycle by increasing carbon flux into and out of this central metabolic pathway. Extended Data Figure 4 summarizes the global importance of wtIDH1 on biochemical pathways, and in particular ones that impact mitochondrial function and redox homeostasis.

Experiments were then pursued to validate wtIDH1 as a bona fide biologic and therapeutic cancer target using both in vitro and in vivo models. In a clonogenic assay, IDH1-knockout had no effect under glucose abundance (e.g., 25 mM, Fig. 2a). However, IDH1−/− cells derived from two different PC cell lines were crippled under glucose withdrawal (Fig. 2a, Extended Data Fig. 5a, b), a consequence of metabolic collapse. N-acetylcysteine (NAC, a glutathione precursor) rescued IDH1−/− cells under glucose withdrawal in these studies, and underscored the role that antioxidant defense plays in IDH1-driven adaptive survival. We expected these findings to translate to animal models, since tumors are nutrient-deprived, relative to well-perfused normal tissues (*7, 11*). A prior study of wtIDH1 knockout mice supports this idea (*28*). Mice were healthy at baseline, but experienced oxidative liver failure injury and death with severe LPS exposure. Herein, IDH1−/− xenografts derived from two separate PC cell lines failed to proliferate like isogenic control xenografts (Extended Data Fig. 5c, d). As with the above-mentioned in vitro studies (Fig. 2a), forced elevation of ambient glucose in the mice, in this case, through the ingestion of sugar water (dextrose water; abbreviated as D30 water), increased intratumoral glucose levels and rescued IDH1−/− xenograft growth (Fig. 2b-d). Increased ambient glucose minimized PC cell dependence on wtIDH1, and had the opposite effect on *IDH1* expression as glucose withdrawal (Fig.1d and Fig. 2e).

**Figure 2.**
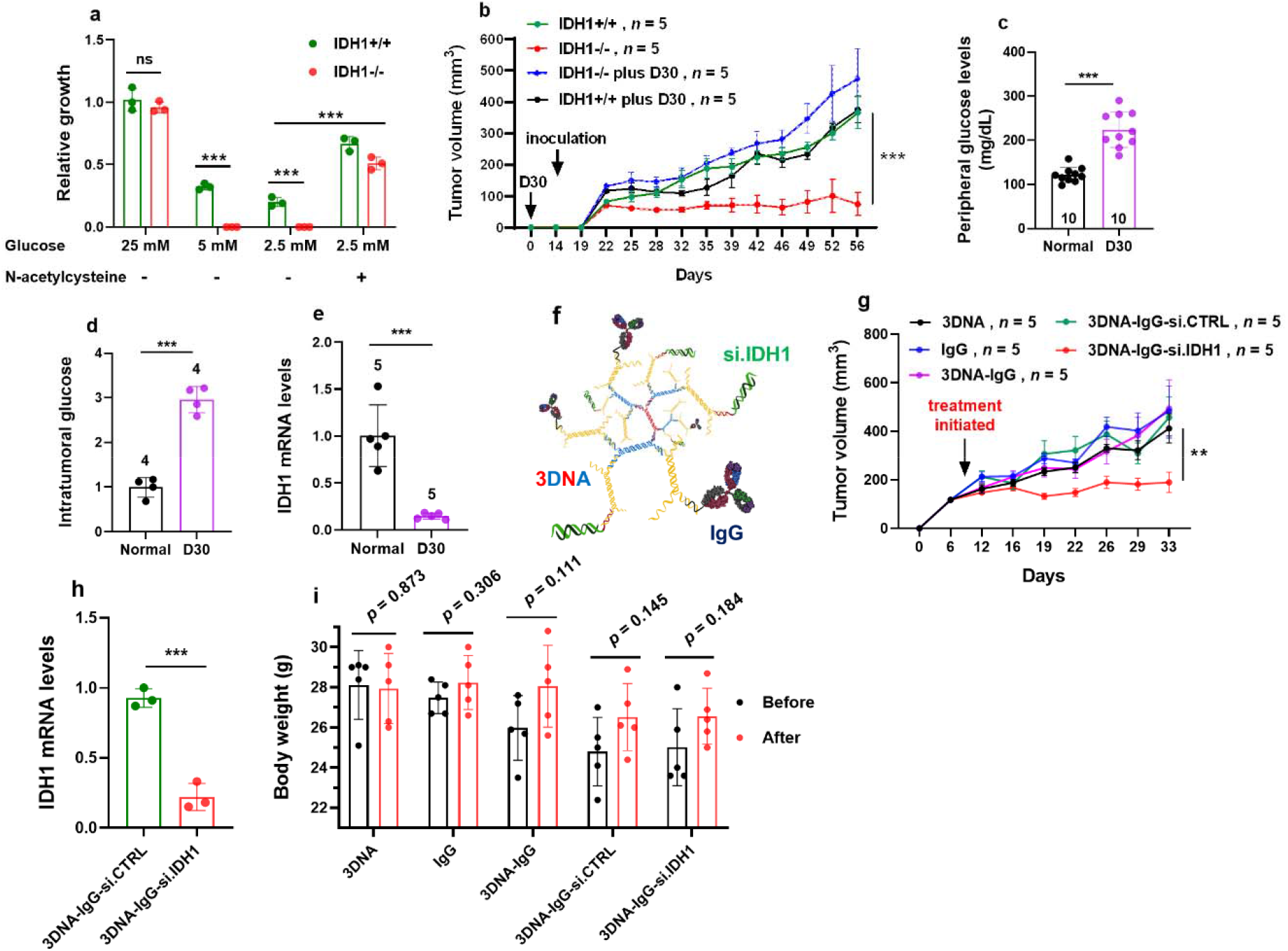
IDH1 is a therapeutic target in pancreatic cancer. **a**, Representative data from a clonogenic growth assay of MiaPaCa-2 PC cells treated with or without a glutathione precursor NAC (1.25 mM) at the indicated conditions. NAC treatment was given 16 hours prior to culturing cells with low glucose conditions; n=3 independent experiments. **b**, Growth of subcutaneous MiaPaCa-2 tumors in mice receiving normal or 30% dextrose water. The number of mice per group is indicated. **c**, Peripheral glucose levels in (**b**). **d**, GC-MS analysis of intratumoral glucose levels (IDH1+/+ vs. IDH1+/+ plus D30 water). **e**, qPCR analysis of *IDH1* transcript normalized to 18S in xenografts shown in (**d**). **f**, Graphical depiction of 3DNA nanocarriers conjugated with si.IDH1 and IgG antibody as a non-specific targeting construct. **g**, Growth of subcutaneous tumors. The number of mice per group is indicated. qPCR analysis of *IDH1* mRNA transcripts (**h**) and body weights (**i**) from indicated treatment arms from experiment (**g**). Data are provided as mean ± s.d. For xenograft growth (**b**, **g**), error bars represent s.e.m. *P* values were calculated using two-tailed, unpaired Student’s *t*-tests. **, P < 0.01; ***, P < 0.001.

In independent mouse experiments, IDH1 was silenced using 3DNA nanocarriers. 3DNA is a nanoscale, biodegradable DNA dendrimer with single-stranded oligonucleotide arms situated at the periphery of the 3DNA structure. These single-stranded oligos can hybridize effector molecules (*29*). In this case, 3DNA was derivatized with siRNAs against the *IDH1* transcript (Fig. 2f). IgG was used as a non-specific targeting moiety, and could potentially be replaced with cancer receptor-specific ligands to improve targeting. Intraperitoneal administration of 3DNA-si.IDH1 significantly slowed xenograft growth in mice, as compared to a variety of control arms (Fig. 2g). IDH1 knockdown in the tumors was validated at the mRNA and protein levels (Fig. 2h, Extended Data Fig. 5e). Mice showed no adverse physical effects in any of the treatment or control arms, and body weights remained stable throughout the experiment (Fig. 2i).

IDH1 therapeutic investigations have principally focused on the development of mutant IDH1 inhibitors (*30*). Less than a handful of reports have focused on wild-type IDH1 inhibition (*21, 30–32*). Both mutant and wild-type isoenzymes possess an allosteric pocket where small molecule inhibitors are believed to compete with positively-charged Mg^2+^ for a negatively charged aspartate residue (Asp^279^) (*31, 33, 34*). The Mg^2+^-bound enzyme is catalytically active, while a low Mg^2+^ state may permit small molecule inhibition at the allosteric site (Extended Data Fig. 6a) (*33*). The Km for Mg^2+^ was previously reported to be roughly 300-fold lower for wtIDH1 than mtIDH1 (20 μM vs. 6 mM, respectively) (*33*). This observation is consistent with the interpretation by researchers that conventional allosteric inhibitors weakly bind wtIDH1 (*30*). They reasoned that Mg^2+^ outcompetes allosteric inhibitors for Asp^279^, even at relatively low Mg^2+^ concentrations, to maintain wild-type enzyme activity (*33, 34*). For this reason, allosteric IDH inhibitors are suspected to have minimal activity against normal tissues (*30*). However, prior studies generally evaluate allosteric IDH1 inhibitors in cell-free systems under conditions that do not represent the tumor microenvironment (*30, 33, 34*). Roughly half of the intracellular magnesium pool is protein-bound, while the remaining fraction exists as free cationic magnesium (*35*). Mg^2+^ levels are dynamic, dependent on antiporter activity, and are lower in stromal rich environments (*35–37*). Thus, the efficacy of allosteric IDH inhibitors on wtIDH1 in tumors remains unknown.

The IDH1 wild-type sequence was first validated in PC, as well as other cancer cell types (Extended Data Fig. 6b). Mg^2+^ levels in serum and tumors were then measured by atomic absorbance spectrophotometry. Consistent with human sera (*38, 39*), circulating Mg^2+^ levels were around 1 mM. Levels in subcutaneous or orthotopic pancreatic tumors were just 20% of these levels. Normal tissues also had relatively low Mg^2+^ levels (Extended Data Fig. 6c). We examined the efficacy of a panel of commercially available mtIDH1 inhibitors against wtIDH1 under standard (similar to serum) and lower (to simulate the tumor microenvironment) Mg^2+^ concentrations. In a cell-based wtIDH1 activity assay, inhibitors were generally ineffective under standard Mg^2+^ concentrations. However, they were all potent at lower Mg^2+^ concentrations (Fig. 3a). In a dose response assay with AG-120 (ivosidenib), drug efficacy improved at lower Mg^2+^ levels, and plateaued below 0.4 mM Mg^2+^ (Extended Data Fig. 6d). This effective concentration is actually higher than the previously reported Km in cell-free assays (*33*). Importantly, this concentration is within the physiologic range observed in PC (Extended Data Fig. 6c).

**Figure 3.**
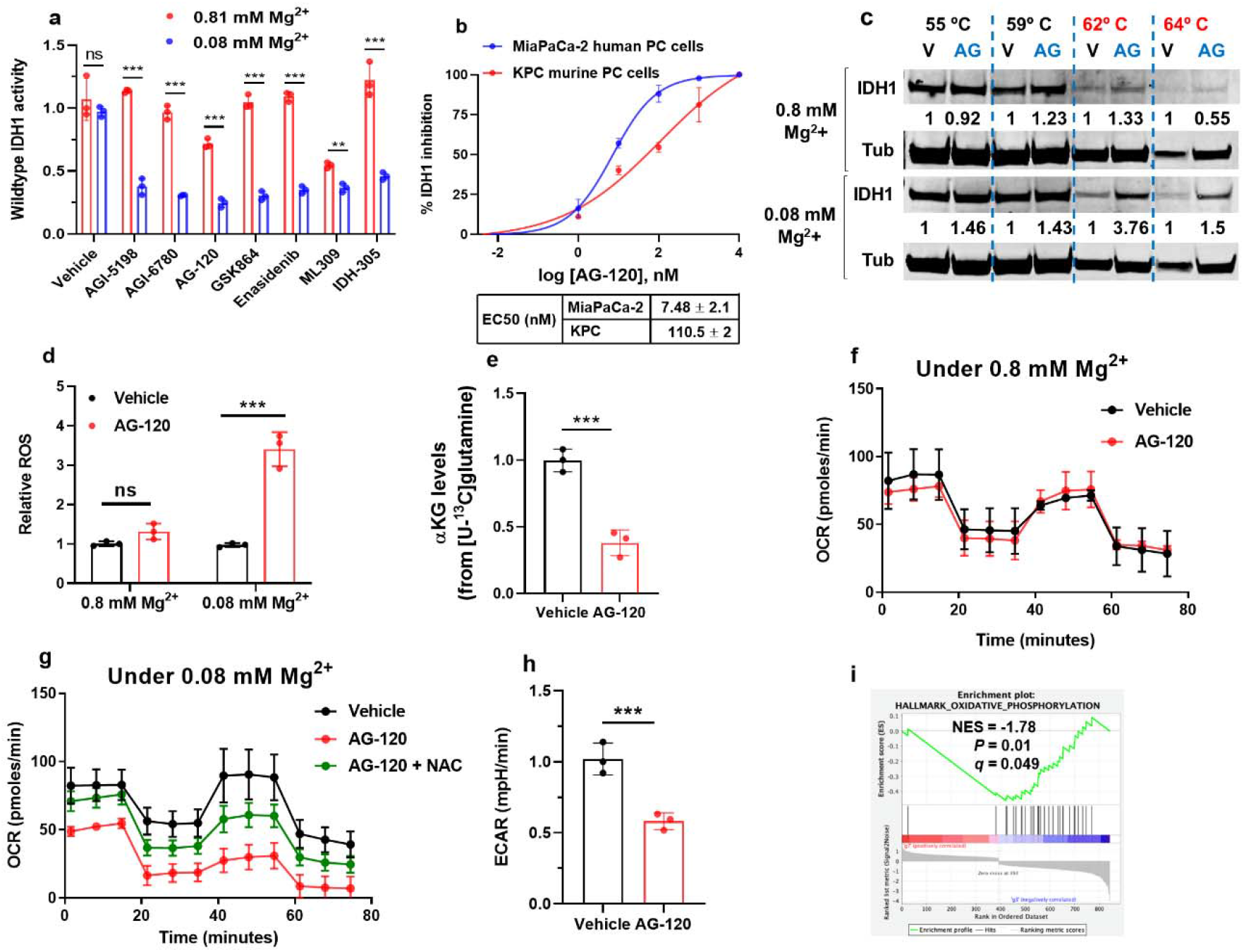
AG-120 is a potent wild-type IDH1 inhibitor under low Mg^2+^ and low glucose. **a**, IDH1 activity analysis in MiaPaCa-2 PC cells first cultured in media containing 0.8 or 0.08 mM MgSO4 and 25 mM glucose for 24h, followed by treatment with a panel of commercial mtIDH1 inhibitors (100 nM) for 6-8 hours; n=2 independent experiments. **b,** IDH1 activity (EC50) of MiaPaCa-2 human PC cells and KPC murine PC cells treated with AG-120 for 24 hours in media containing 0.08 mM MgSO4; n=2 independent experiments. **c**, Western blot analysis to evaluate thermal stability of the IDH1 protein in MiaPaCa-2 PC cells treated with vehicle or AG-120 (1 μM) in media containing 0.8 or 0.08 mM MgSO4 for 6-8 hours. **d**, ROS levels were detected by DCFDA assay in MiaPaCa-2 PC cells treated with AG-120 (1 μM) for 24 hours under the indicated conditions, and in media containing 2.5 mM glucose. **e**, GC-MS analysis of αKG abundance (all isotopologues) in MiaPaCa-2 PC cells cultured in 2.5 mM glucose for 24 hours; n=2 independent experiments. Representative oxygen consumption rate in MiaPaCa-2 PC cells treated with vehicle or AG-120 in media containing 0.8 Mg^2+^ mM and 2.5 mM glucose (**f**), treated with vehicle, AG-120 or AG-120 plus NAC in media containing 0.08 mM Mg^2+^ and 2.5 mM glucose (**g**) for 30 hours. NAC treatment was given 16 hours prior to AG-120 treatment. **h**, Extracellular acidification rates (ECAR) in MiaPaCa-2 PC cells cultured in media containing 0.08 mM Mg^2+^ and 2.5 mM glucose for 30 hours. **i**, GSEA of oxidative phosphorylation genes in MiaPaCa-2 PC cells treated with vehicle or AG-120 (10 μM) under 0.08 mM Mg^2+^ and 2.5 mM glucose for 48 hours. Data are provided as mean ± s.d. from three independent experiments, unless indicated. *P* values were calculated using two-tailed, unpaired Student’s *t*-tests. **, P < 0.01; ***, P < 0.001.

AG-120 is orally available and well-tolerated. The drug is currently FDA-approved at a dose of 500 mg daily to treat medically refractory, IDH1-mutant acute myeloid leukemia (AML) (*40*). Additionally, AG-120 has shown activity against IDH1-mutant cholangiocarcinoma in a phase III trial (*41*). We performed a series of in vitro and in vivo experiments with AG-120 to understand the drug’s effect on wild-type IDH1 cancers, particularly under physiologically relevant conditions. In a cell-free wtIDH1 activity assay, the drug was 350-fold more potent at 0.3 mM Mg^2+^, as compared to 10 mM (Extended Data Fig. 6e). This was replicated in cell-based assays. In MiaPaCa-2 PC cells cultured media containing 0.08 mM Mg^2+^ and 25 mM glucose, the estimated EC50 after 24h of AG-120 was 7.48 nM (Fig. 3b). The drug was also potent, albeit to a lesser degree, against murine wtIDH1 (Fig. 3b), which has 95.7% sequence homology to the human form. Of note, the compound had no inhibitory activity against IDH2, which is the mitochondrial homolog of IDH1 (Extended Data Fig. 6f). A cellular thermal shift assay provided indirect evidence of binding between AG-120 and wtIDH1 (Fig. 3c). At higher temperatures, the compound stabilized the protein under low Mg^2+^ concentrations, while the effect was not observed at higher Mg^2+^ levels. Of note, low Mg^2+^ alone (in the absence of AG-120), had a negligible effect on PC cells (Extended Data Fig. 6g). AG-120 treatment caused a surge in ROS under low Mg^2+^ and low glucose conditions, yet no such effect was observed in the presence of higher Mg^2+^ levels (Fig. 3d). A similar pattern was apparent in the nucleus, as AG-120 induced substantial oxidative damage to DNA under low Mg^2+^ and low glucose (Extended Data Fig. 7a).

Stable isotope tracing experiments mimicked the results observed in IDH1−/− cells. Incubation with [U-^13^C]glutamine resulted in a sharp decline in ^13^C-labeled αKG under low glucose conditions (Fig. 3e, similar to Extended Data Fig. 3j). AG-120 had no effect on oxygen consumption in PC cells cultured under standard Mg^2+^ concentrations (Fig. 3f). On the other hand, the drug severely impaired oxygen consumption when PC cells were cultured in low glucose and low Mg^2+^. An antioxidant, NAC, partially restored mitochondrial function under these conditions (Fig. 3g, similar to Fig. 1i and Extended Fig Data. 2f). AG-120 also reduced ECAR under low glucose and Mg^2+^ concentrations (Fig. 3h).

Transcriptomic studies of PC cells treated with AG-120 resulted in 861 differentially expressed genes (FDR < 0.05) (Extended Data Fig. 7b). Remarkably, oxidative phosphorylation-related genes were among the top significantly downregulated genes observed by GSEA analysis (normalized enrichment score of –1.78, *p* = 0.01, *q* < 0.05) (Fig. 3i, Extended Data Fig. 7c). A focused heatmap of enriched genes in the oxidative phosphorylation pathway indicates a high proportion of these genes were significantly downregulated with AG-120 treatment (Extend Data Fig. 7d). Interestingly, *IDH1* was one of these genes. These data do not indicate if transcript suppression was a direct result of AG-120 treatment or rather a second level consequence of mitochondrial injury, the known mechanism of AG-120 suggests the latter explanation is more likely.

Thus, two conditions are required for AG-120 efficacy against wtIDH1 tumors: 1) low Mg^2+^ (to inhibit wtIDH1 activity), and 2) low nutrients (such that wtIDH1 is required for cell survival). If either condition is absent, the drug is ineffective as a wtIDH1 inhibitor, as shown in murine and human PC cell culture models (Fig. 4a and Extended Data Fig. 7e, respectively). Extended Data Figure 7f summarizes these principles. Since both conditions are only present in tumors to our knowledge, wtIDH1 inhibitors are particularly attractive as therapeutics. Whereas chemotherapy becomes less effective under nutrient-deprivation (*4*), wtIDH1 inhibition becomes even more effective under these conditions (Extended Data Fig. 7g).

**Figure 4.**
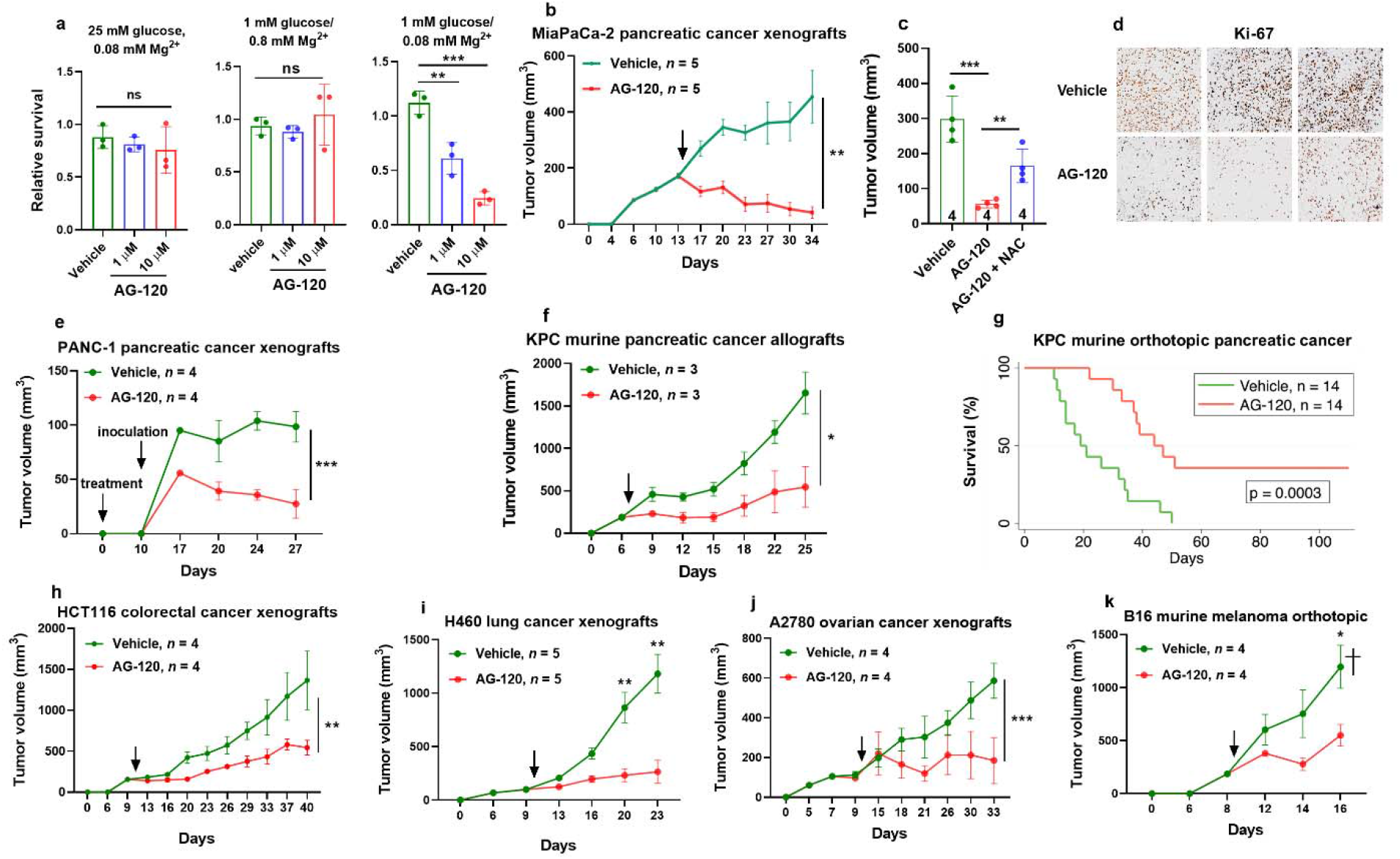
AG-120 inhibits wtIDH1 cancer growth in mice. **a**, Relative clonogenic growth of MiaPaCa-2 PC cells treated with vehicle or AG-120 under different levels of glucose and Mg^2+^ for 72 hours; n=3 independent experiments. For all animal experiments, mice were treated orally with vehicle or AG-120 (150 mg/kg). Treatment started when tumors were palpable unless indicated, as shown in the graphs with an arrow. The number of mice per group is indicated. **b**, MiaPaCa-2 xenograft growth in nude mice treated with vehicle or AG-120 for the indicated days. **c**, Independent MiaPaCa-2 xenograft experiment in nude mice. In the indicated arm, N-acetylcysteine (NAC, 1.2 g/L) was administered in the drinking water for three days, followed by vehicle, AG-120, or AG-120 plus NAC. **d**, Ki-67 staining (20X) of three xenografts treated with vehicle or AG-120 for seven days. **e**, Mice were initially treated orally with AG-120 for nine days. On day 10, PANC-1 PC cells were subcutaneously implanted into nude mice. Tumor growth was monitored for 17 days post cell inoculation. **f,** Growth of subcutaneous xenografts derived from murine PC (KPC cells) transplanted into nude mice. Mice were treated with vehicle or AG-120. **g**, Survival analysis of murine PC orthotopically transplanted into the pancreas of immunocompetent mice. Mice were treated with vehicle or AG-120. **h-k**, Growth of HCT116 (colon), H460 (lung), A2780 (ovarian) in nude mice, and B16 (murine melanoma) xenografts in immunocompetent mice, treated with vehicle or AG-120 for the indicated time interval. † in (**k**) indicates that one mouse in the vehicle group was found dead with a destroyed tumor; the penultimate recorded tumor volume was used for the analysis. Data are provided as mean ± s.d. For xenograft growth (**b**, **e-f**, **h-k**), error bars represent s.e.m. *P* values were calculated using two-tailed, unpaired Student’s *t*-tests. Median survival was analyzed using the Kaplan-Meier estimate and compared by the log-rank test. *, P < 0.05; **, P < 0.01; ***, P < 0.001.

WtIDH1 tumors were transplanted into mice to test in vivo efficacy of AG-120. The dosage was identical to the dose used previously to treat mtIDH1 tumors (150 mg/kg (*42*)). Human tumors were engrafted into nude mice, while murine cancers were engrafted into immunocompetent mice (C57BL/6J), unless indicated. Mice tolerated the drug well and maintained their weights over time (Extended Data Fig. 7i). AG-120 proved effective as a single agent against every single wtIDH1 cancer tested, including pancreatic cancer (Fig. 4b-g), colorectal cancer (Fig. 4h), lung cancer (Fig. 4i), ovarian cancer (Fig. 4j), and melanoma (Fig. 4k). In each of these instances, the anti-tumor effect of AG-120 was comparable or superior (i.e., tumor shrinkage) to a previous study of anti-IDH1-mutant tumor therapy (*43*). Similar to in vitro experiments (Extended Data Fig. 5a, b), NAC (1.2 g/L water) partially rescued PC growth (Fig. 4c), showing in an animal model that AG-120 disrupts redox balance in tumor cells. In the immunocompetent orthotopic PC model, AG-120 improved median survival by more than two-fold (19 vs. 44 days) (Fig. 4g).

These data offer novel insights. First, wtIDH1 represents a true metabolic vulnerability in cancer cells and is a bona fide therapeutic target across a wide range of wtIDH1 cancers. Second, the products of wtIDH1 oxidative decarboxylation (NADPH and αKG) support antioxidant defense and mitochondrial function. Both biologic processes are required for PC survival under austere conditions present in the tumor microenvironment. Third, mutant-IDH1 inhibitors, including FDA-approved AG-120, are potent wild-type IDH1 inhibitors under low Mg^2+^ and low glucose conditions. Since pancreatic and other tumors share this feature, these drugs are compelling investigational agents for these expanded indications. We believe that this study provides a strong rationale for human trials testing this hypothesis in patients.

---

**Extended Data Fig. 1.**
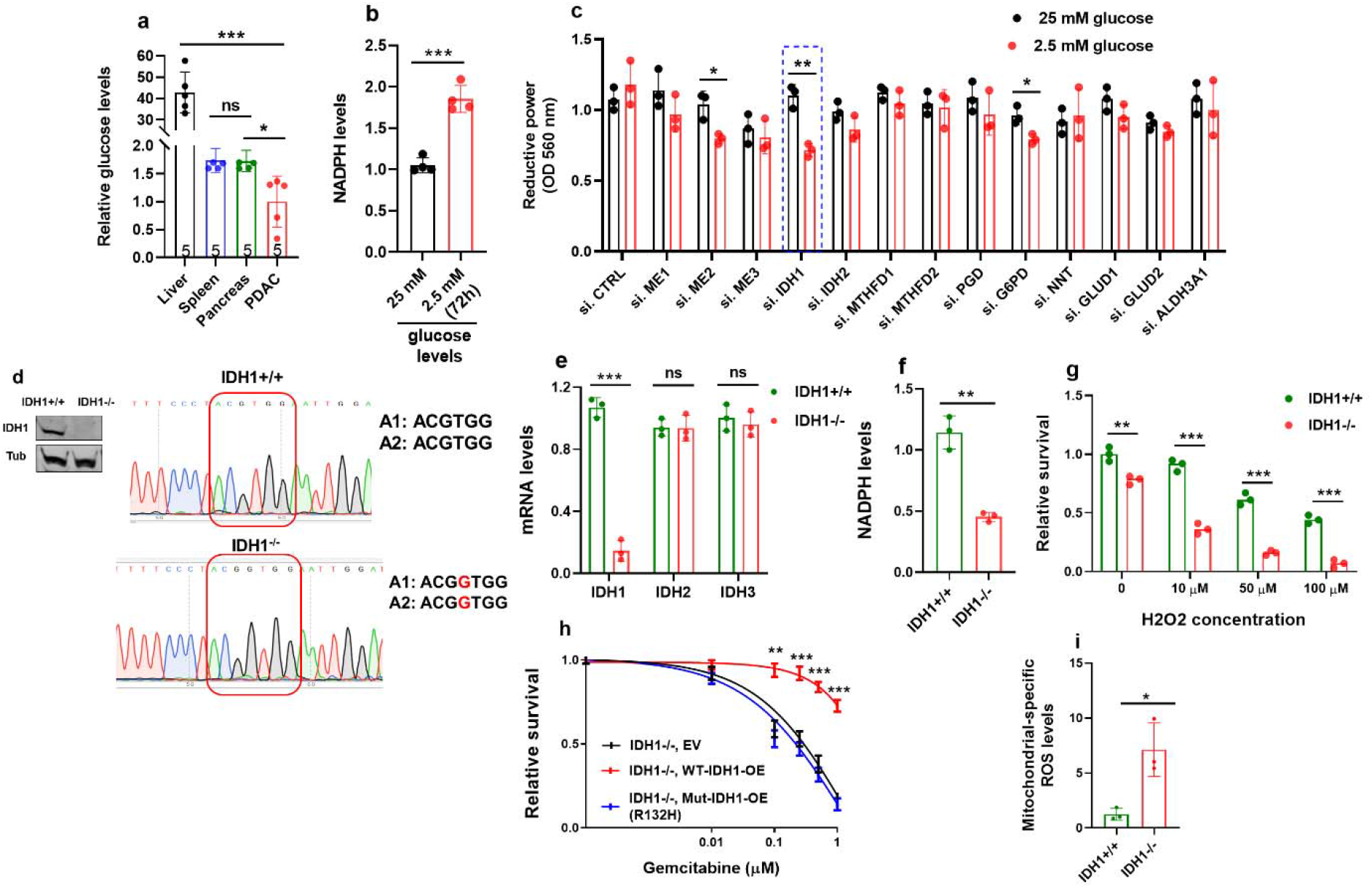
IDH1 loss enhances sensitivity to oxidative stress. **a**, Relative glucose levels in liver, spleen, pancreas and subcutaneous MiaPaCa-2 PC xenografts from nude mice. The number of mice analyzed per group is indicated. **b**, NADPH levels in PANC-1 PC cells under the indicated conditions for 72 hours, n=4 independent experiments. **c**, Reductive power, as measured by an MTT assay, after transient transfection of siRNAs against NADPH-generating enzymes in PANC-1 PC cells. Cells were incubated under the indicated conditions for 72 hours. **d**, Western blot and sanger sequencing validation of IDH1 in IDH1+/+ and IDH1−/− MiaPaCa-2 PC cells. **e**, mRNA expression quantitation in MiaPaCa-2 PC cells by qPCR of IDH1, IDH2 and IDH3 transcripts. mRNA was normalized to 18S. **f**, NADPH levels in MiaPaCa-2 PC incubated under 2.5 mM glucose for 72 hours. **g**, Relative survival as detected by Trypan blue assay in Hs766T cells. Cells were treated with either vehicle or hydrogen peroxide, and incubated with 2% serum for 72 hours. **h**, Relative survival as detected by PicoGreen assay in IDH1−/− MiaPaCa-2 PC cells, transiently transfected with empty vector (EV), plasmid overexpressing catalytically active, or altered (R132H) IDH1, and treated with gemcitabine for 5 days under 5 mM glucose (relative glucose withdrawal). **i**, Mitochondrial-specific ROS measured in MiaPaCa-2 PC cells, under 2.5 mM glucose for 72 hours. Data are provided as mean ± s.d. from three independent experiments, unless indicated. *P* values were calculated using two-tailed, unpaired Student’s *t*-tests. *, P < 0.05; **, P < 0.01; ***, P < 0.001.

**Extended Data Fig. 2.**
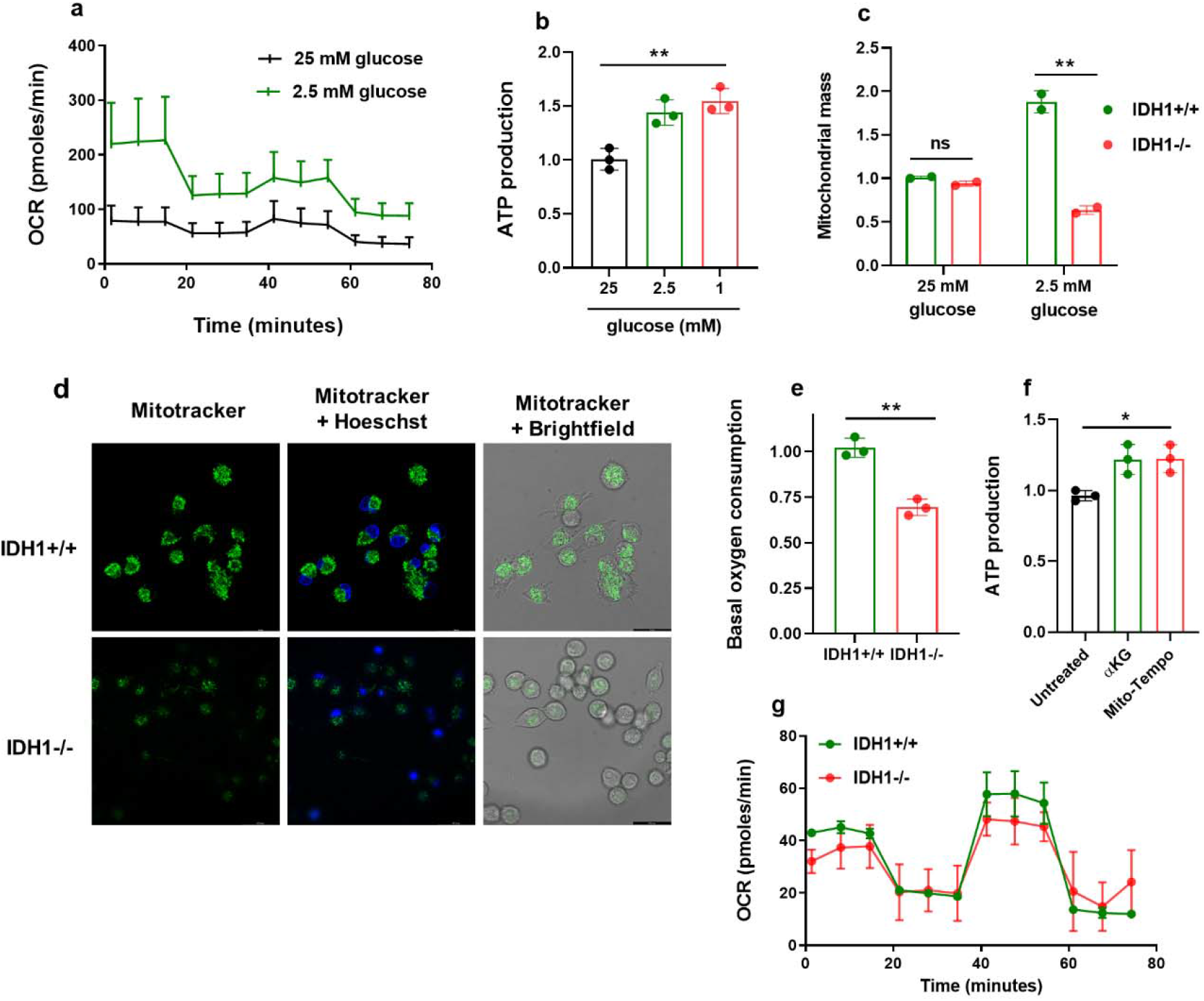
IDH1−/− cells have impaired mitochondrial function. **a,** Representative oxygen consumption rates in MiaPaCa-2 PC cells cultured under the indicated glucose levels for 30 hours. **b**, ATP levels in MiaPaCa-2 PC cells cultured under the indicated conditions for 24 hours. **c**, Mitochondrial mass in MiaPaCa-2 PC cells cultured under the indicated conditions for 48 hours. **d**, Confocal microscopy images of MiaPaCa-2 PC cells in an independent experiment cultured in 1 mM glucose for 48 hours. **e**, Basal OCR in Hs766T PC cells cultured in 1 mM glucose for 24 hours; n=2 independent experiments. **f**, ATP levels in MiaPaCa-2 PC IDH1−/− cells cultured with αKG (4 mM) and Mito-Tempo (100 μM) for 24 hours. **g**, OCR in MiaPaCa-2 PC cells cultured in 25 mM glucose for 30 hours; n=2 independent experiments. Data are provided as mean ± s.d. from three independent experiments, unless indicated. *P* values were calculated using two-tailed, unpaired Student’s *t*-tests. *, P < 0.05; **, P < 0.01.

**Extended Data Fig. 3.**
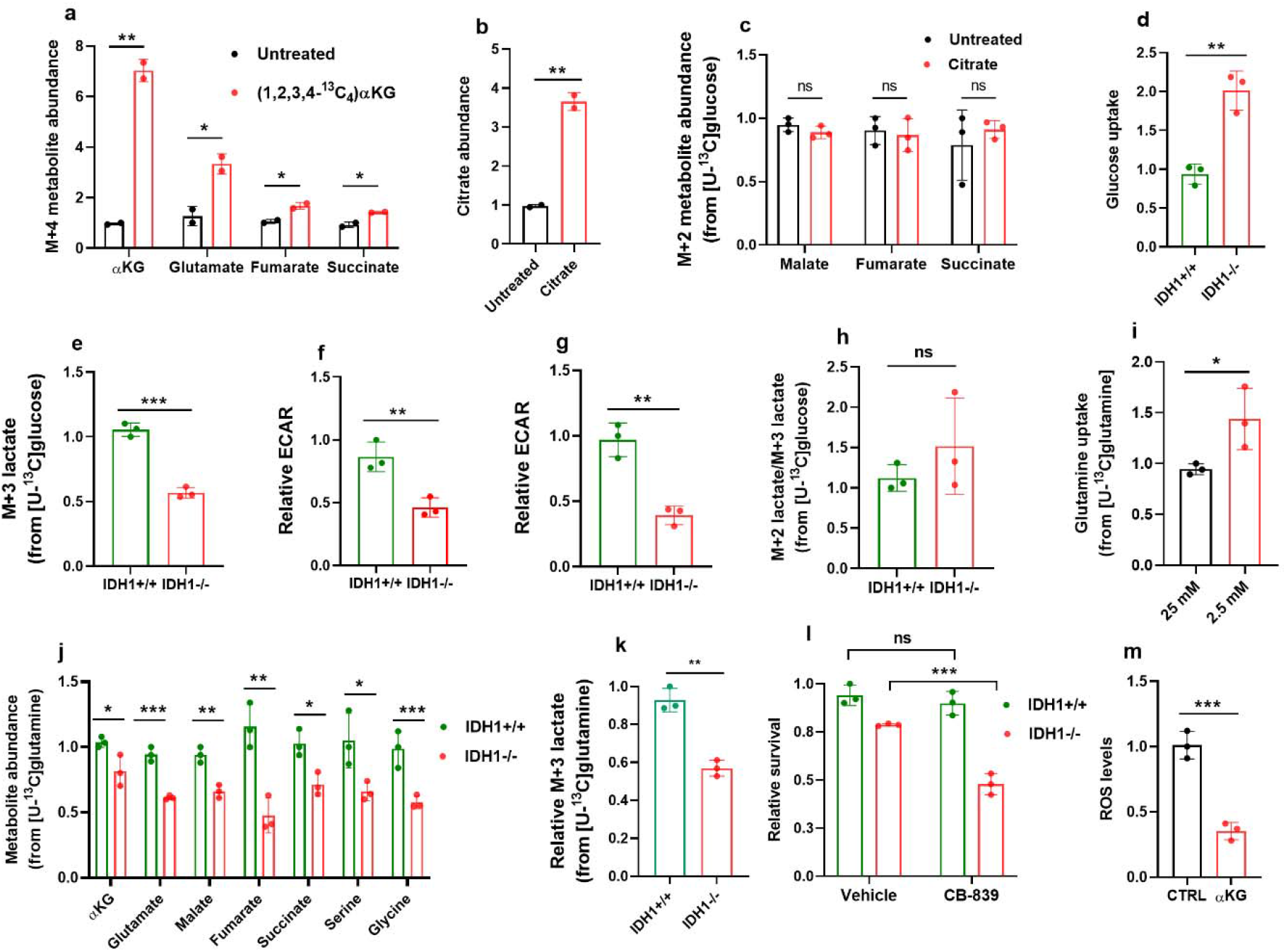
IDH1-directed metabolic reprogramming under nutrient withdrawal. **a**, GC-MS analysis of M+4 metabolite enrichment in MiaPaCa-2 IDH1−/− cells, cultured in media containing (1,2,3,4-^13^C_4_)αKG (4 mM) for 10 hours; n=2 independent experiments. **b,** Sodium citrate (4 mM) was added to the cell culture media. IDH1−/− cells were incubated for 10 hours; n=2 independent experiments. **c**, MiaPaCa-2 PC IDH1−/− cells were cultured in media containing [U-^13^C]glucose (2.5 mM) and sodium citrate (4 mM) for 10 hours; n=2 independent experiments. **d**, Glucose uptake in MiaPaCa-2 PC cells, cultured in 2.5 mM glucose for 30 hours. **e**, Representative m+3 lactate levels from MiaPaCa-2 PC cells cultured in 2.5 mM glucose. Relative glycolytic rates measured in MiaPaCa-2 (**f**) and Hs766T (**g**) PC cells under low glucose conditions for 24 hour. **h**, M+2 lactate/ M+3 lactate ratios derived from [U-^13^C]glucose, in MiaPaCa-2 PC IDH1−/− and +/+ cells cultured in 2.5 mM glucose for 24 hours. **i,** Relative glutamine uptake was quantified via mass spectrometry. MiaPaCa-2 PC cells were cultured in 25 or 2.5 mM glucose for 24 hours. **j**, Metabolite abundance (all isotopologues) in MiaPaCa-2 PC cells was quantified. **k**, Representative M+3 lactate levels from MiaPaCa-2 PC cells cultured in 2.5 mM glucose. **l**, Survival of isogenic MiaPaCa-2 PC cells cultured in 2.5 mM glucose for 24 hours, followed by CB-839 (1 μM) treatment for an additional 24 hours. **m**, ROS levels in MiaPaCa-2 PC IDH1−/− cells, cultured with αKG (4 mM), and in 2.5 mM glucose for 48 hours. Data are provided as mean ± s.d. from three independent experiments, unless indicated. *P* values were calculated using two-tailed, unpaired Student’s *t*-tests. *, P < 0.05; **, P < 0.01; ***, P < 0.001.

**Extended Data Fig. 4.**
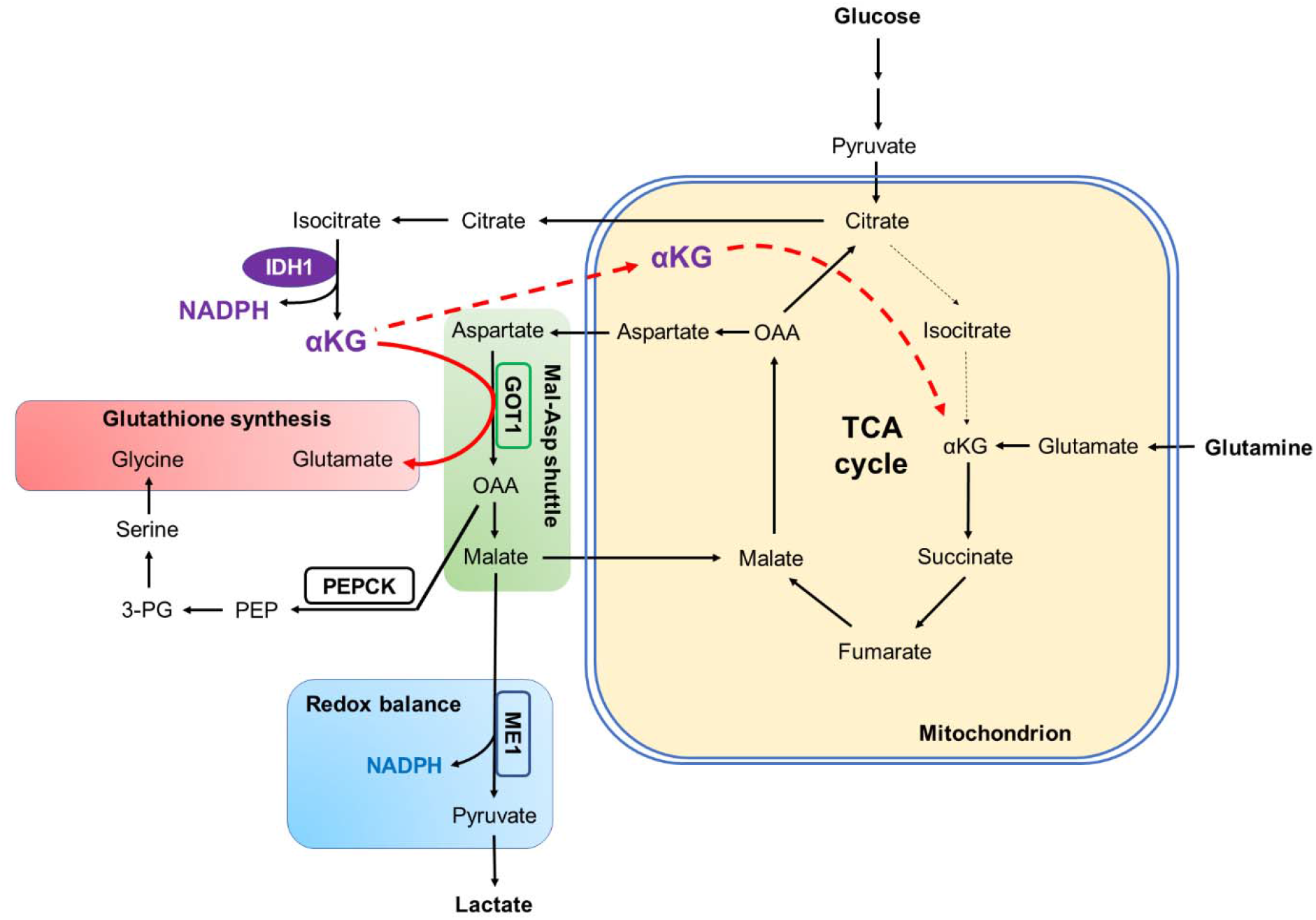
Schematic overview of IDH1-mediated glucose and glutamine metabolism in PC cells under nutrient withdrawal. IDH1-derived αKG drives carbon flux through the TCA cycle, and influences other interconnected pathways. OAA, oxaloacetate; PEP, phosphoenolpyruvate; PEPCK, phosphoenolpyruvate carboxykinase; and 3-PG, 3-phosphoglycerate.

**Extended Data Fig. 5.**
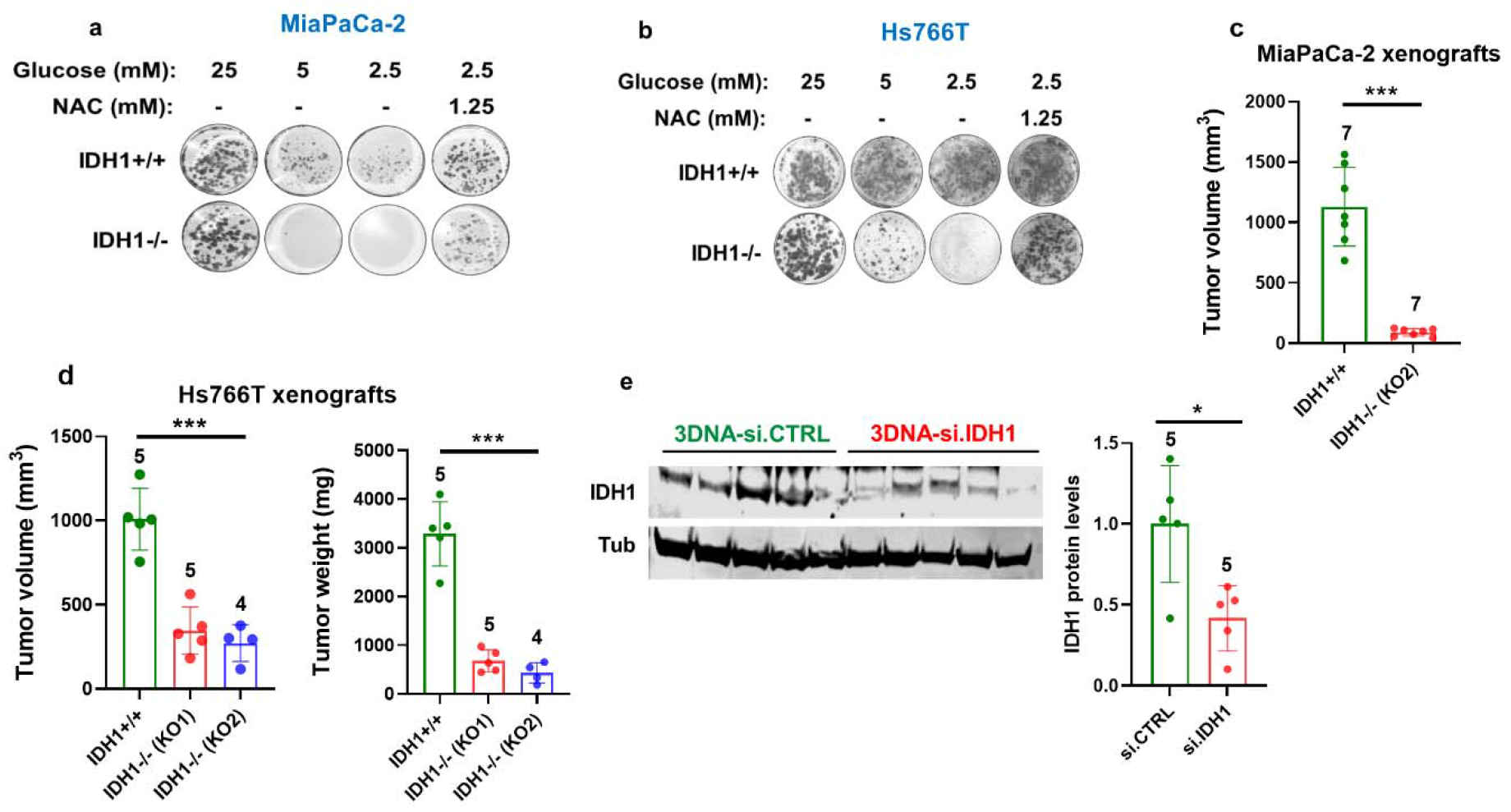
Reduced growth in IDH1-knockout PC cells and xenografts. **a,** Representative wells from the clonogenic growth assay graphed in Fig. 2a. **b**, Clonogenic growth of Hs766T PC cells cultured under the indicated conditions. **c-d**, Growth of subcutaneous tumors from nude mice. **e**, Western blot analysis of IDH1 protein in xenografts treated with 3DNA-si.CTRL or 3DNA-si.IDH1 graphed in Fig. 2b. Data are provided as mean ± s.d. from three independent experiments, unless indicated. *P* values were calculated using two-tailed, unpaired Student’s *t*-tests. *, P < 0.05; ***, P < 0.001.

**Extended Data Fig. 6.**
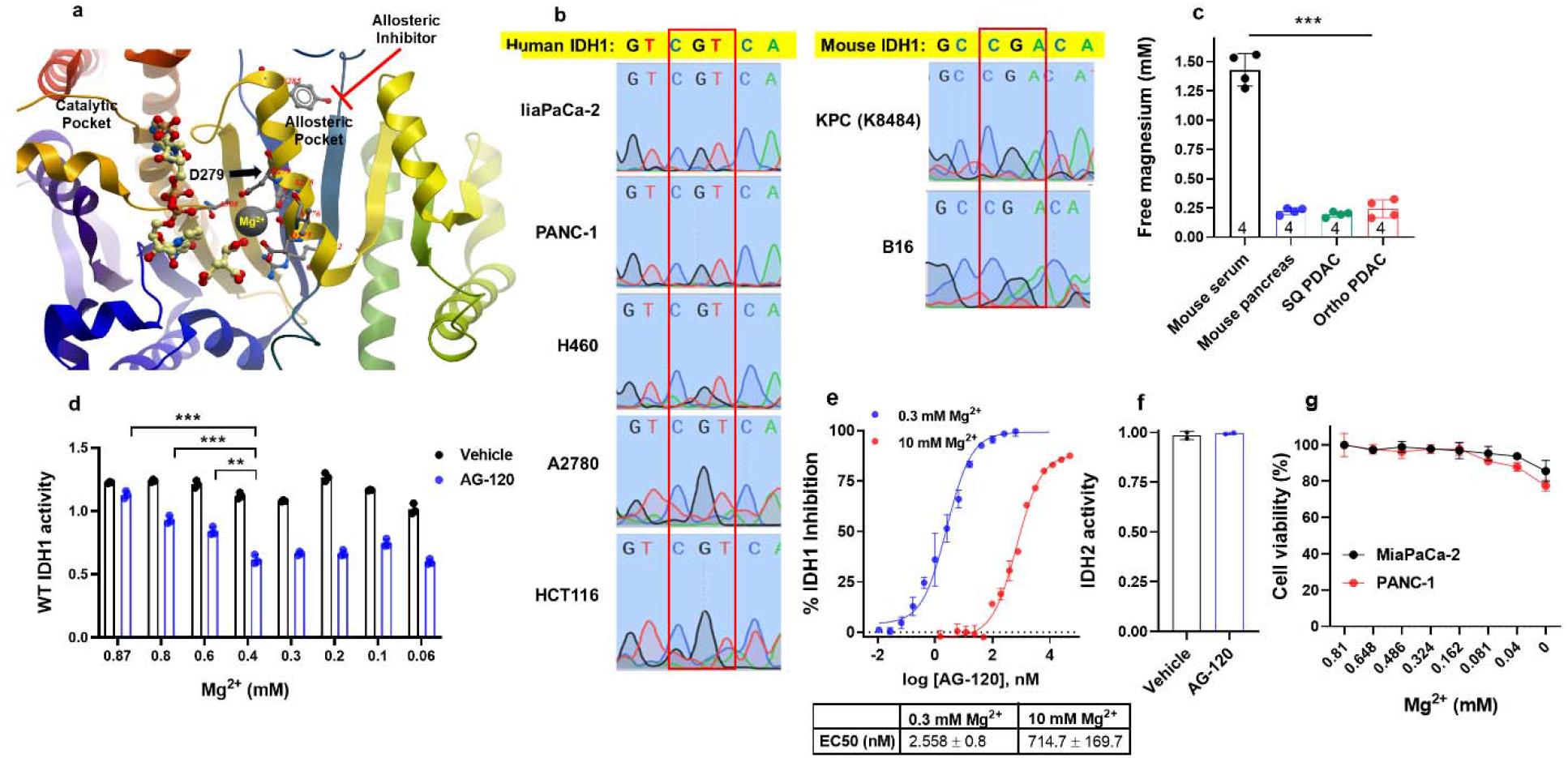
Mg^2+^ levels influence the efficacy of allosteric IDH inhibitors against the wtIDH1 enzyme. **a**, Wild-type IDH1 from the Protein Database. Mg^2+^ is believed to interact with D279 (Asp279) in the allosteric pocket (accession number: PDB 1t0l). Under high Mg^2+^ conditions, the allosteric inhibitor is ineffective. Under low Mg^+2^ conditions, the drug is effective. **b**, Sanger sequencing of amplicons correlating with codon 132 of the wtIDH1 gene. The reference wildtype sequence is shown. **c**, Free magnesium levels in MiaPaCa-2 xenografts and orthotopic KPC tumors vs. serum. Levels in normal tissues are also shown. The number of analyzed tissues is indicated. **d**, IDH1 activity measurements (cell-based assay) in MiaPaCa-2 PC cells; n=2 independent experiments. **e**, IDH1 activity (cell-free assay) upon treatment with AG-120 under the indicated conditions; n=2 independent experiments. **f**, AG-120 effects on IDH2 activity (cell-based assay) in MiaPaCa-2 PC cells; n=2 independent experiments. **g**, Cell viability of PC cells, cultured in MgSO4 at the indicated concentrations for five days. Data are provided as mean ± s.d. from three independent experiments, unless indicated. *P* values were calculated using two-tailed, unpaired Student’s *t*-tests. **, P < 0.01; ***, P < 0.001.

**Extended Data Fig. 7.**
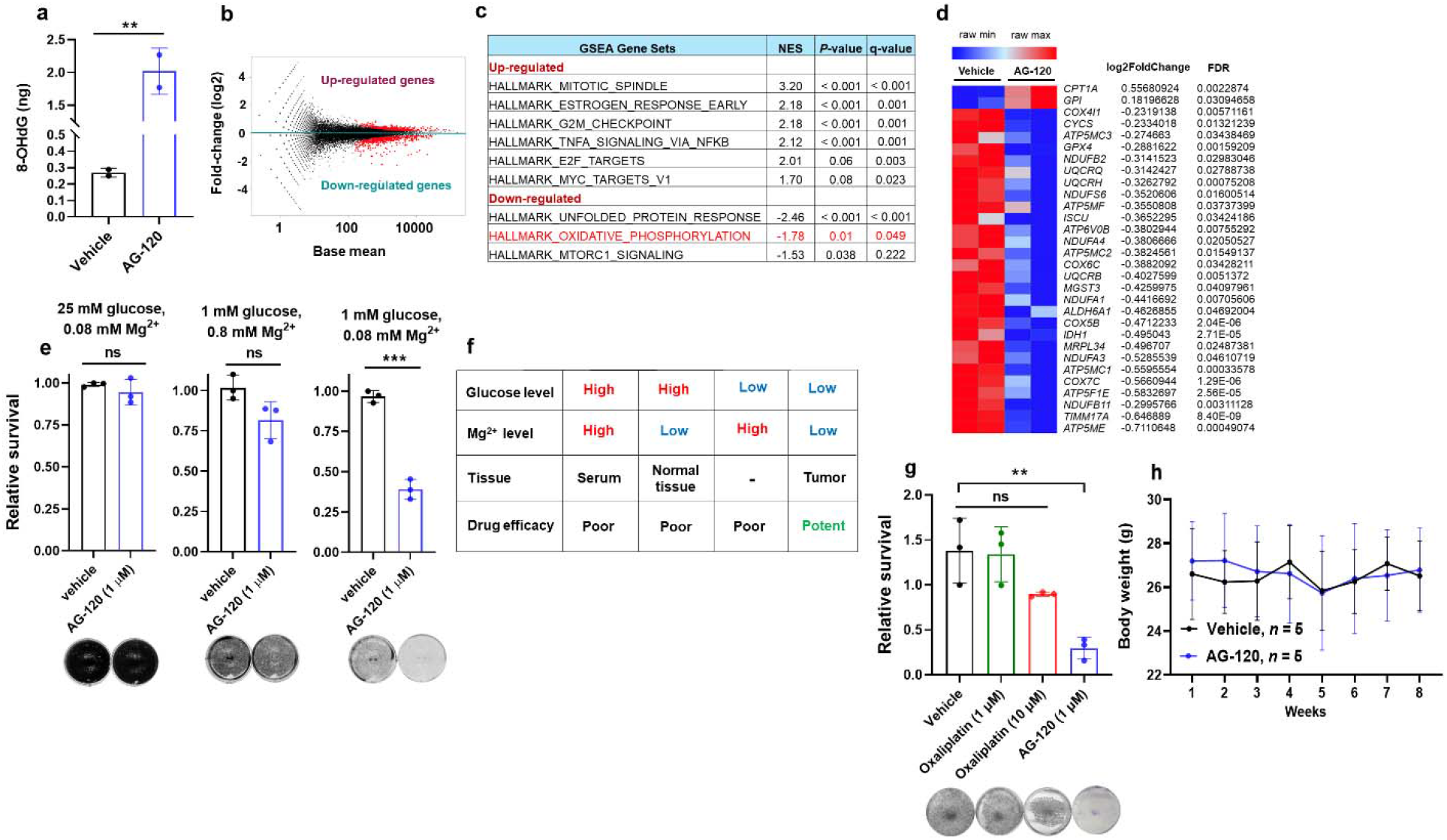
Diverse effects of AG-120 against cancer cells under low glucose and low Mg^2+^ conditions. **a**, Oxidized DNA in MiaPaCa-2 PC cells detected by 8-hydroxy-deoxyguanosine (8-OHdG) levels, under 2.5 mM glucose for 48 hours; n=2 independent experiments. MA plot representing differential gene expression (**b**) and GSEA analysis (**c**) in MiaPaCa-2 PC cells after AG-120 (10 μM) treatment (vs. vehicle) under 2.5 mM glucose for 48 hours; n=2 independent replicates. **d**, Heatmap of differentially expressed genes associated with oxidative phosphorylation under the indicated conditions; n=2 independent replicates. **e**, Relative clonogenic growth of KPC cells (murine PC cells, K8484) treated with vehicle or AG-120 under the indicated conditions for 72 hours. **f**, An overview of key factors impacting the efficacy of allosteric IDH1 inhibitors against wtIDH1 cancer cells. **g**, Relative clonogenic growth of KPC cells cultured under 1 mM glucose and 0.08 mM Mg^2+^ for 72 hours, and treated as indicated. **i**, Body weights of mice treated with AG-120 at 150 mg/Kg, or vehicle. Data are provided as mean ± s.d. from three independent experiments, unless indicated. For (**f**), error bars represent s.e.m. *P* values were calculated using two-tailed, unpaired Student’s *t*-tests. **, P < 0.01; ***, P < 0.001.

## Acknowledgments

We would like to thank Dr. Sanford Markowitz for his valuable insights and support; Dr. Alpaslan Tasdogan for his helpful suggestions with the manuscript; Dr. Darren Carpizo for generously providing K8484 KPC cells; Dr. Andrea Romani for technical help with magnesium measurements; Dr. William Schiemann for providing luciferase expressing plasmid; Richard Lee for imaging assistance; and CWRU Genomic core facility for sequencing assistance. We acknowledge the use of the Leica TCS SP8 confocal microscopy in the CWRU SOM Light Microscopy Imaging core, made available through funding from NIH ORIP S10OD024996. Grant support for this research comes from American Cancer Society MRSG-14-019-01-CDD (J.M.W), Postdoctoral Fellowship grant NIH/NCI F32CA247466 (A.V), American Cancer Society 134170-MBG-19-174-01-MBG (J.M.W.), NIH/NCI R37CA227865-01A1 (J.M.W), the Case Comprehensive Cancer Center GI SPORE 5P50CA150964-08 (J.M.W), and University Hospitals research start-up package (J.M.W.) We are grateful for additional support that contributed to this work from the John and Peggy Garson Family research fund, The Jerome A. and Joy Weinberger Family research fund, and Mark Levine in memory of Ethel Levine.

## Author Contributions

A.V., J.R.B., H.B., J.S. and J.M.W. designed the study and interpreted experiments. A.V. performed most of the experiments and composed the original draft of this manuscript. A.V., J.J.H., H.J.G., J.C. and K.D. performed molecular and cell biology experiments. A.V., J.B., R.G. and J.M.W. performed 3DNA experimental design and provided reagents. A.V. and O.H. performed magnesium detection. A.V., E.P., V.C. and I.B. performed metabolomics, isotope tracing experiments and analysis. H.J.G provided reagents, helped with animal studies protocol, animal ordering and mycoplasma testing. I.K., J.J.H., and H.J.G helped A.V. to perform in vivo experiments. A.A. performed transcriptomic analysis. F.S.M provided IDH1 inhibitors. L.R. helped design, interpret and oversee animal experiments testing melanoma cells. M.Z. and R.W. helped with data interpretation. A.V. and J.M.W. primarily composed the manuscript. All authors contributed to manuscript revisions.

## Methods

### Cell lines and cultures

All cell lines were obtained from ATCC, except murine PC cells (KPC K8484: Kras^G12D^; Trp53^R172H/+^; Pdx1-Cre). Routine mycoplasma screening was performed. Cell lines were maintained at 37° C and 5% CO2. For standard cell culture, cells were grown in DMEM containing 25 mM glucose and 4 mM glutamine, and supplemented with 10% FBS, 1% penicillin/streptomycin, and prophylactic doses of plasmocin (Life Technologies, MPP-01-03) to prevent mycolplasma infection. Glucose withdrawal was performed to simulate low glucose conditions in the PDAC microenvironment. For these experiments, glucose-free DMEM (Life Technologies, 21013-024) was supplemented with 2.5 mM glucose (Sigma), 10% FBS, and penicillin-streptomycin. For experiments with low magnesium, glucose-, glutamine- and magnesium sulphate-depleted DMEM (Cell Culture Technologies, 964DME-0619) was supplemented with the indicated concentrations of MgSO4, FBS, and antibiotics. Nutrients were added, when indicated, for each experiment. In this manuscript, standard or high Mg^2+^ refers to levels that approximate serum (0.8 to 1.5 mM); low levels approximate the tumor microenvironment and are generally below 0.4 mM.

### CRISPR/Cas9 editing of IDH1 in pancreatic cancer cells

IDH1 knockout was performed using a gRNA: GTAGATCCAATTCCACGTAGGG (Sigma, target ID, HS0000323225). Plasmid transfections were performed with lipofectamine 2000 (Life technology, 11668-027). Negative control plasmid (CRISPR06-1EA) was used in isogenic cells. After 48h, EGFP-expressing cells were sorted by FACS. Clones were expanded and genomic DNA extracted for Western blot verification of IDH1 deletion. Cells with IDH1 deletion and isogenic controls are referred to as IDH1−/− and IDH1+/+, respectively.

### Cell viability assays

Cell viability was measured by cell counting using Trypan blue reagent (Life Technologies, 15250061) or by DNA quantitation via PicoGreen dsDNA assay (Life Technologies, P7589).

### Clonogenic assay

1000-2000 cells per well were plated in a 6-well plate. The media was not changed during experiments unless indicated. Upon completion of the experiments, colonies were fixed in reagent containing 80% methanol and stained with 0.5% crystal violet. To determine relative growth, dye was extracted from stained colonies with 10% acetic acid and the associated absorbance measured at 600 nm. With regard to AG-120 experiments, cells were first cultured with low MgSO4 media (0.08 mM) for 24 hours, followed by AG-120 treatment under 2 mM glutamine, 5% FBS and indicated glucose levels for 72 hours.

### ROS and 8-OHdG quantification

Cells were incubated in a 96-well plate with 10 μM H2-DCFDA (Life Technology, D399) for 45 min in serum-free media to detect total intracellular ROS. For mitochondrial-specific ROS, cells were incubated with 5 μM MitoSOX Red (Life Technologies, M36008) for 30 min in serum-free media. Cells were washed with PBS, and fluorescence measured according to manufacturer instructions using a GloMax Explorer system (Promega). 8-OHdG was measured according to the manufacturer’s instructions (Abcam, AB201). Readouts were normalized to cell number.

### Quantitative RT-PCR

Total RNA was extracted using PureLink RNA isolation (Life Technologies, 12183025) and treated with DNase (Life Technologies, AM2222). cDNA was synthesized using 1 μg of total RNA, oligo-dT and MMLV HP reverse transcriptase (Applied Biosystems, 4387406). All PCR reactions were performed in triplicate, and primer sequences are provided in the supplementary section.

### RNA-sequencing and analyses

RNA quality was assessed via the Agilent 2100 Bioanalyzer (Agilent Technologies). Strand-specific RNA-seq library was prepared using NEBNext Ultra II Directional RNA Library Prep Kit (NEB, Ipswich, MA) according to the manufacturer’s protocols. RNA-sequencing was performed using 150-bp paired-end format on a NovaSeq 6000 (Illumina) sequencer. RNA-seq quality was checked by running FastQC, and TrimGalore was used for adapter and quality trimming. RNA-seq reads were mapped against hg38 using STAR (v 2.7.0e) aligner with default parameters. DESeq2 analysis with a FDR value <0.05 was used to generate a list of differentially expressed genes. Gene set enrichment analysis was performed using GSEA (v 4.1.0) with Hallmark gene sets (v 7.2).

### DNA sequencing

DNA from human and murine PC cells were isolated using the DNeasy blood and tissue kit (Qiagen) according to the manufacturer’s protocol. A portion of the IDH1 gene exon 4 containing the Arg132 was amplified using two pairs of primers (for human IDH1, F: 5′-ACCAAATGGCACCATACGA-3′; R: 5′-TTCATACCTTGCTTAATGGGTGT-3′; for mouse IDH1, F: 5′-ATTCTGGGTGGCACTGTCTT-3′; R: 5′-CTCTCTTAAGGGTGTAGATGCC-3′), and sequenced using one of the amplification primers. Gene segments corresponding to the targeted regions were amplified to validate knockout efficacy (F: 5′-GAGGGCTAGCTCAGAAAC-3′, R: 5′-CATTGTACTATTCTTAGCCACTG-3′). Sanger sequencing was performed using following primer: 5′-TGGCGGTTCTGTGGTAGAG-3′. PCR reactions were carried out as follows: 95 °C for 30 sec, 60 °C for 30 sec, and 72 °C for 40 sec for a total of 40 cycles. The products were purified, and Sanger sequencing was performed using one of the amplification primers.

### Small RNA interference

siRNA transfections were performed using Lipofectamine 2000. siRNA oligos were obtained from Life technology. Gene knockdown validation was determined 72 hours after siRNA transfections via qPCR.

### Western blot analysis

Total protein was extracted with RIPA buffer (Pierce, 89900) supplemented with protease inhibitor (Life Technologies, 1861280) and quantified using the BCA Protein Assay (Thermo Scientific). Proteins were separated on 4-12% Bis-Tris gels (Life Technologies, MW04125) and transferred to PVDF membranes. Membranes were probed with antibodies against IDH1 (Invitrogen, GT1521) overnight at 4 °C and alpha-Tubulin (Invitrogen, 11224-1-AP) for 1-2 hours at room temperature. Blots was probed with secondary antibodies customized for the Odyssey Imaging system.

### Cellular Thermal Shift assay

Thermal shift experiments were performed as previously described (*44*). Briefly, cells were cultured in media containing the indicated concentrations of MgSO4 under standard condition (25 mM glucose, 4 mM glutamine and 10% FBS) for 24 hours. Then, cells were treated with either vehicle (DMSO) or AG-120 (1 μM) for 6-8 hours under indicated Mg^2+^ levels. Cells were then trypsinized, washed with PBS, suspended in PBS (containing protease inhibitors), lysed through three freeze-thaw cycles at liquid nitrogen-37 °C, and then heated at the indicated temperatures for three minutes. Preparations were spun down at 13,000 rpm for 10 min at 4 °C to remove denatured proteins, and supernatants were loaded on a Western blot gel. Western blots were performed as described.

### IDH activity assay

Cell-based IDH activity is measured through the quantitation of IDH-dependent NADPH synthesis (Abcam, ab 102528). The reaction measures the activity of both IDH1 and IDH2 isoenzymes. IDH1 activity is specifically measured using the same assay after transient siRNA silencing of IDH2. The opposite approach is used to estimate IDH2 activity. All readouts were normalized to cell number. Cell-free WT IDH1 activity was performed by a fluorometric assay measuring the NADPH levels at 355/460 nm. Assays contained 0.25 nM human recombinant IDH1 (RD systems), 30 μM isocitrate, and 30 μM NADP^+^ in 20 μL of 100 mM Tris-HCL, pH8, 0.2 mM DTT, 0.05% CHAPS, and the indicated MgCl2 levels. Formation of NADPH was measured over 40 min at 5 min intervals.

### Cell bioenergetic and glucose uptake assays

Oxygen consumption rates (OCR) and extracellular acidification rates (ECAR) were measured using the Seahorse XFp Extracellular Flux Analyzer (Seahorse Bioscience) according to the manufacturer’s instructions. For OCR experiments, cells were cultured in 4 mM glutamine, 5% FBS, at indicated glucose levels, for 24-36 hours. With regard to AG-120 experiments, cells were treated with AG-120 (2 μM) under 4 mM glutamine, 5% FBS, and indicated glucose levels. NADPH levels (Abcam, ab65349), ATP (Abcam, 83355) and glucose uptake (Sigma, MAK083) were performed according to the manufacturer’s instructions. All readouts were normalized to cell number.

### Confocal live imaging

Confocal laser scanning microscopy was performed on a Leica TCS SP8 (Leica Buffalo Grove, IL) using a 63x/1.4 NA oil immersion objective. Nuclear counterstaining was performed using Hoeschst dye with excitation and emission wavelengths of 405nm and 430-470nm, respectively. Light detection was performed using a photomultiplier tube detector. MitoTracker Green was imaged at excitation and emission wavelengths of 488nm and 500-520nm using a hybrid detector. Channels were acquired sequentially.

### Magnesium measurement of tissues in mice

Once animals were euthanized, tissues were collected, washed with 250 mM sucrose and quickly snap frozen in liquid nitrogen. 50 mg of tissues were homogenized in 10% sucrose via sonication on ice, followed by centrifugation at 12,000 rpm for 10 min at 4 °C. Supernatants were removed and examined for free Mg^2+^ content by atomic absorbance spectrophotometry (AAS; Agilent Technologies) upon appropriate dilution. Readouts were normalized according to homogenate volume and protein content.

### Metabolic profiling

Cells were grown in complete growth media in T25 flasks. Experiments were performed in biological triplicates. For metabolic profiling of IDH1−/− and +/+ cells, processing began when cells reached 50% confluency. Standard culture media was changed to the low glucose media (2.5 mM glucose, 4 mM glutamine supplemented with 10% dialyzed FBS) for 38-hour incubation. Subsequently, cells were incubated in the media containing [U-^13^C]glucose (Cambridge Isotope Laboratories, CLM-1396-1) or [U-^13^C]glutamine (Cambridge Isotope Laboratories, CLM-1822-H-0.1) for 10 hours. For rescue experiments, cells were incubated after adding either unlabeled αKG (4 mM) or sodium citrate (4 mM) to media containing stable isotope tracers. For experiments testing the effect of AG-120 treatment, media was additionally modified to contain low magnesium concentrations (0.08 mM MgSO_4_) for 24 hours when indicated, followed by AG-120 (1 μM) treatment for 2 hours. Media was then exchanged with low glucose media as indicated, and for 18 hours. Cells were allowed to equilibrate for three hours in the media containing either [U-^13^C]glutamine. Cells ready for processing were placed on ice, washed 3X with cold PBS, and lysed using in ice-cold 80:20 methanol:water. For animal tissues and xenograft tumors, fragments weighing 15-25 mg were homogenized in ice-cold 80:20 methanol:water using a sonicator at 4 °C. Unlabeled _γ_-hydroxybutyrate (50 μM) was added to homogenized lysate as an internal standard. Cell extracts were centrifuged at 14,000 rpm for 15 min at 4°C. Supernatants were then dried using nitrogen gas. Samples were then incubated with 30 μl of freshly made MOX solution (40 mg methoxyamine hydrochloride dissolved in 1ml of pyridine) for 45 min at 45°C to protect keto groups, followed by the incubation with 70 μl of MTBSTFA (N-(tert-butyldimethylsilyl)-N-methyltrifluoroacetamide) for 45 min at 45°C to derivatize metabolites. For glucose measurements, after sample drying with nitrogen gas, dried lysates were mixed with pyridine:acetic anhydride (1:2) solution and incubated at 60°C for 30 min. Samples were allowed to air dry at room temperature, and then reconstituted in 100 μl of ethyl acetate. GC-MS analyses were performed using a Hewlett Packard 5973 Turbo Pump Mass Selective Detector and a Hewlett Packard 6980 Gas Chromatograph equipped with a DB-5ms GC Column (60 m x 0.25 mm x 0.25 um, Agilent Technologies). Data were processed with Agilent Enhanced Data Analysis Software. Samples were injected in splitless mode. The column temperature was initially set at 100°C and held for one minute, then ramped 8°C/min until 170°C and held for 5 min. Samples were then ramped 5°C/min until 200°C, and held for 5 min. Finally, samples were ramped 10°C/min until 300°C, and held 10 min. Masses were monitored via the Scan acquisition mode. Metabolite counts were further normalized using cell number from a parallel culture flask, and plated at an equivalent cell density to reflect equal sample loading.

### In vivo studies

All experiments involving mice were approved by the CWRU Institutional Animal Care Regulations and Use Committee (IACUC, protocol 2018-0063). Six-to-eight week-old, female, athymic nude mice (Foxn1 nu/nu) were purchased from Harlan Laboratories (6903M) through the ARC CWRU. For flank xenograft experiments, 1×10^6^ cells were suspended in 200 μL of a PBS:matrigel solution (1:1) and injected subcutaneously into the right flank of mice. For orthotopic experiments, 1×10^4^ Luciferase-expressing KPC K8484 cells were suspended in 50 μL of a PBS:matrigel solution (1:1) and injected into the pancreas of C57BL/6 mice at 12 weeks of age. Equal number of male and female mice were used for survival analysis. Mice were anesthetized using isoflurane gas. After ensuring appropriate depth of anesthesia, a 0.5 cm left subcostal incision was made to gain access into the peritoneal cavity. The tail of the pancreas was externalized and the mixture was carefully injected into the pancreas. The pancreatic tail was left alone for one minute to limit any tumor cell leakage, and then returned to the peritoneal cavity. The incision was then closed with a combination of permanent suture and skin clips. On postoperative day 6-7, the presence of pancreatic tumors was confirmed with bioluminescence imaging, IVIS, using 100 μl intraperitoneal Luciferin injection (50 mg/mL). Mice with confirmed tumors were then randomized to the indicated treatment regimens. For the indicated experiments, hyperglycemia was induced by allowing mice to drink high glucose content water (30% dextrose; abbreviated as D30 water) starting two weeks before cancer cell implantation. 3DNA nanocarriers were prepared as described previously (*45*). Briefly, reagents (containing 0.0625 μg/μL 3DNA, 0.057 μg/μL IgG, 0.758 μM si.RNA) were diluted in PBS and injected intraperitoneally every 3 days for the indicated period of time. For the indicated experiments, AG-120 (Asta Tech, 40817) was suspended in 10% PEG-400, 4% Tween-80 (3% for MiaPaCa-2 and PANC-1 xenografts) and 86% saline for maximal dissolution of drug. After tumors became palpable (0.6 cm in diameter or 120-150 mm^3^), the cocktail was administrated twice daily at 150 mg/kg, with at least 8 hours between doses. An equal volume of vehicle was administrated to control mice. For the PANC-1 xenograft experiment, treatment was initiated prior to tumor injection. NAC water (1.2 g/L) was refreshed twice weekly for the indicated experiments. For all flank xenograft experiments, tumor volumes were measured twice per week using a caliper (volume= lengthxwidth^2^/2). Body weights were measured once weekly for the indicated time.

